# Characterization of an *Agarophyton chilense* oleoresin containing PPARγ natural ligands with insulin-sensitizing effects in a C57BL/6J mouse model of diet-induced obesity and antioxidant activity *in Caenorhabditis elegans*

**DOI:** 10.1101/2021.03.16.435726

**Authors:** Claudio Pinto, María Raquel Ibañez, Gloria Loyola, Luisa León, Yasmin Salvatore, Carla Gonzalez, Victor Barraza, Francisco Castañeda, Rebeca Aldunate, Loretto Contreras-Porcia, Karen Fuenzalida, Francisca C Bronfman

## Abstract

The biomedical potential of the edible red seaweed *Agarophyton chilense* (former *Gracilaria chilensis*) has not been explored. Red seaweeds are enriched in polyunsaturated fatty acids and eicosanoids, which are known natural ligands of the PPARγ nuclear receptor. PPARγ is the molecular target of thiazolidinediones (TZDs), drugs used as insulin sensitizers to treat type 2 diabetes mellitus. TZDs medical use is limited due to undesired side effects, the fact that it has triggered the search for partial agonists without the TZDs side effects.

We produced *A.chilense* oleoresin (*Gracilex®*) that induces the activation of PPARγ without inducing adipocyte differentiation, similar to PPARγ partial agonists. In a model of high-fat diet-induced obesity in male mice, we showed that treatment with *Gracilex®* improves insulin sensitivity, normalizing altered glucose and insulin parameters. *Gracilex®* was enriched in palmitic acid, arachidonic acid, oleic acid, and lipophilic antioxidants such as ß-carotene and tocopherols. *Gracilex®* possesses antioxidant activity in vitro (CUPRAC) and increased the antioxidant capacity *in vivo* in *Caenorhabditis elegans.* These findings support the idea that *Gracilex®* is an excellent source of healthy lipids targeting PPARγ with antioxidant activity and a high nutraceutical value.

## Introduction

The edible Chilean red macroalgae *Agarophyton chilense* (formerly *Gracilaria chilensis*) (1), commonly known as “pelillo”, has been used as a food and medicinal herb since pre-Hispanic times in Chile, as indicated by findings at the archaeological site of Monte Verde (~14,000 years ago) (2). *A. chilense* is distributed in New Zealand (including Chatham Island), and South America (3–6), and although this species has good nutritional potential it is mainly exploited for the extraction of the hydrocolloid agar-agar (7, 8).

Seaweeds contain numerous molecules with nutritional and biomedical potential since they are rich in vitamins, minerals, essential trace elements, polysaccharides, and dietary fiber and are a good source of protein and lipids compared to terrestrial plants (9). Seaweed pigments such as carotenoids and xanthophylls have excellent nutraceutical value and show metabolic effects in humans (10). Red algae such as *A. chilense*, are an excellent source of healthy essential fatty acids such as polyunsaturated fatty acids (PUFAs), including eicosapentaenoic acid (EPA), arachidonic acid (ARA), and oleic, linoleic, and alpha-linolenic fatty acids. Other well-studied phytochemicals include fucoidans, polyphenols, tocopherols, and phytosterols. The reported bioactivities of seaweed, including antidiabetic, anti-inflammatory, antioxidant and anti-neurodegenerative activities, are varied (11, 12). Thus, in recent years, there has been increased interest in identifying novel bioactive compounds that demonstrate health benefits to confirm and increase the added value to edible red seaweeds (8, 13).

Various methods of extraction are used to obtain enriched fractions derived from seaweeds containing antioxidant and pigment molecules (14, 15), proteins (16), polysaccharides (17) and lipids (18, 19). Particularly in the *Gracilaria* genus, lipid extractions obtained by classical chloroform:methanol mixtures have revealed a complex lipid composition of glycerolipids and omega-3 and omega-6 polyunsaturated fatty acids and their oxidized derivatives, known as oxylipins (18–21). Oxylipins are synthesized either by the metabolic action of specific lipoxygenases (LOXs) or by nonenzymatic reactions on PUFAs such as EPA, ARA, linoleic acid and linolenic acid (22–24). Interestingly, the oxylipin structure is similar or equivalent to that of mammalian prostaglandins, leukotrienes, and eicosanoids (25); which are well-known ligands of peroxisome proliferator-activated receptors (PPARs) (26–28).

Lipid components of seaweeds (e.g., tocopherols, xanthophylls, carotenoids, phytosterols, PUFAs, and oxylipins) target members of the nuclear superfamily of ligand-inducible transcription factors (NRs), including peroxisome proliferator-activated receptors (PPARs), retinoid X receptors (RXRs), and the liver X receptor (LXR), which together regulate the homeostasis of cholesterol, fatty acids, glucose, insulin sensitivity and inflammation (29, 30). These studies suggest that the benefit of algae-derived food or nutraceuticals is due to their multiple targets, including NRs.

The PPARs NR subfamily present three receptor subtypes (PPARa, PPARß/∂, and PPARγ) encoded by distinct genes, and they are obligate heterodimers of the RXR receptors (31). PPARs have attracted increased attention as therapeutic targets given the physiological relevance of the processes in which they are involved. These processes include lipid and glucose homeostasis, promotion of thermogenesis, lipoprotein synthesis, reduction of triglycerides, regulation of glucose levels in the blood, and the generation and maintenance of fat tissue (31–34). Additionally, PPAR activation decreases inflammatory processes by repressing proinflammatory genes regulated by NF-kappaB (35). Finally, these molecules contribute to reducing ROS (reactive oxygen species) by increasing the expression of antioxidant enzymes and increasing mitochondrial biogenesis (36). The three PPARs have natural ligands, including PUFAs and eicosanoids (37–41). All of these lipids activate PPARs with low affinity (micromolar range) compared to high-affinity synthetic agonists such as fibrates and thiazolidinediones (TZDs). In particular, fibrates, drugs used to treat dyslipidemia, target PPARa, favoring fatty acid beta-oxidation, increasing HDL, and lowering triglyceride levels (34, 42). PPARγ has been the best-studied subtype because it is the molecular target of thiazolidinediones (TZDs), which act as full agonists of PPARγ. These drugs were designed in the late 1990s for the treatment of type 2 diabetes mellitus (T2DM). PPARγ is a crucial regulator of blood sugar levels, improving insulin sensitivity in patients and animal models of T2DM (43). However, in early 2007, adverse effects of these drugs, including weight gain, fluid retention, osteoporosis, heart failure, and hepatotoxicity, among others, were described (43–45). These side effects have led to restricted access in the United States and a recommendation for removal from the market in Europe and other jurisdictions (46). At the molecular level, TZDs such as rosiglitazone (RGZ) stabilize the ligand-binding site of PPARγ in an “on” or “off” state. In contrast, partial agonists generate various dynamic and slow conformational changes by recruiting different coactivators and corepressors compared to TZDs. Of note, some partial agonists increase insulin sensitivity without adipogenic effects, such as those induced by TZDs (32,33,47,48). For instance, the partial agonist or selective PPARγ modulator (SPPARM) INT131 has shown improvements in insulin sensitization and glycemic control in T2DM patients with minor side effects compared with those observed by treatment with the full PPARγ agonist pioglitazone (49). In addition to synthetic SPPARMs, natural bioactive compounds with weak or partial PPARγ affinity have shown similar properties and are effective in reducing insulin resistance (50). For example, in a rat model of diet-induced obesity, amorfrutins, a diet-derived ligand for PPARγ, improved insulin sensitization and inflammatory and metabolic parameters (51).

T2DM is a chronic endocrine disease characterized by hyperglycemia in the blood and resistance to the action of insulin, leading to severe neurological and cardiovascular lesions, including myocardial infarction, coronary heart disease, and nonalcoholic fatty liver disease, among others. Although there are genetic risk factors, obesity is considered to explain 60-80% of diabetes cases (52–54).

There are various classes of oral drugs for glycemic management, such as biguanides (metformin), sulfonylureas (SUs), and TZDs. Currently, the first-line oral antihyperglycemic is metformin. When glucose resistance begins, metformin is used in conjunction with the patient’s lifestyle as treatment (55). After metformin, SUs and TZDs are widely used, although they can cause adverse effects (see above) (56).

The majority of patients with T2DM show a deterioration in blood glucose levels over time that cannot be controlled with metformin alone. The escalation dose for metformin and SUs is limited, so patients must incorporate a second-line drug to maintain glycemic control (56, 57). Recently, the use of nutraceuticals such as berberine extracts and omega-3 fatty acids has been suggested for glycemic management and dyslipidemias associated with T2DM (58). Experts have emphasized the use of nutraceuticals and functional foods to treat or mitigate T2DM due to the advantages of having a mix of molecules with simultaneous effects. The Asian continent is familiar with medicine derived from natural products. Therefore, interest in the use of nutraceuticals, including products derived from the sea, such as seaweeds (9), for the treatment of T2DM has increased (59).

Despite the rich diversity of lipids and unique phytochemical composition of red macroalgae lipids, the biomedical potential of lipids derived from *A. chilense* has not been addressed. Therefore, we started our studies by evaluating the ability of an oleoresin derived from *A. chilense (Gracilex®)* to activate PPARγ. We also studied the effect of *A. chilense-*derived lipids on adipocyte differentiation and their capacity to improve the metabolic dysfunction induced by a high-fat diet (HFD). We further characterized the fatty acid profile and the antioxidant capacity of *Gracilex® in vitro* using a copper ion reducing antioxidant capacity (CUPRAC) assay and *in vivo* using a *Caenorhabditis elegans* model of resistance to oxidative stress. Our results indicated that *A. chilense* lipids contain PPARγ activators that act as partial agonists since they do not induce adipocyte differentiation like RGZ. Consistently, the treatment of obese male mice (HFD) reduced glucose and insulin levels increased by HFD, without increasing adiponectin, the hormone that regulates the growth of fat tissue (60). *Gracilex®* possesses an antioxidant capacity *in vitro* and *in vivo* that correlates with high levels of tocopherols and ß-carotene. Our study is the first to examine whether lipids derived from *A. chilense* have nutraceutical or biomedical applications, emphasizing the idea that *A. chilense* can be used as a source of other molecules in addition to its value as a source of agar-agar. More studies will be required to determine the nature of the bioactive molecules present in *Gracilex®*, with particular emphasis on those that have the potential to act as SPPARM-like ligands and that show strong antioxidant activity.

## 2. Material and Methods

### 2.1 Reagents

HPLC grade dichloromethane, cyclohexane and water were purchased from Merck (Darmstad, Germany). Cell culture reagents such us DMEM (Dulbeccós Modifed Eagle Medium), RPMI 1640, opti-PRO™, GlutaMAX^TM^ I supplement, Tripsin-EDTA 0.5% and penicillin/streptomycin were purchase from Gibco BRL (LIFE Technologies Inc, Gaithersburg, USA). Fetal bovine serum and horse serum were obtained from HyClone Laboratories,Inc (Thermo Fisher Scientific, MA, USA) and Lipofectamine^TM^ 2000 from Gibco BRL (LIFE Technologies, MA, USA). Drugs and chemicals, such us FMOC-Leu, MTT (3-(4,5-dimethylthiazol-2-yl)-2,5-diphenyl tetrazolium bromide), ONPG (*ortho*-Nitrophenyl-β-galactoside), NAC (N-acetylcysteine), cell culture grade dimethyl sulfoxide (DMSO), NP-40, Oil red O, dexamethasone, isobutylmethylxanthine (IBMX) and insulin were purchase in Sigma Chemical Co. (MO, USA). Rosiglitazone and T0070907 from Cayman Chemicals (Ann Arbor, MI, USA) and INT131was obtained from MolPort (NY, USA). Plasmids used to measure the transcriptional activity of PPARγ (see below) were a donation of R. M. Evanśs laboratory (Howard Hughes Medical Institute, Gene Expression Laboratory, The Salk Institute for Biological Studies, La Jolla, CA, USA).

### 2.2 Sampling of biomass

Vegetative *A. chilense* individuals, approximately sixty centimeters in length, were collected from two areas along the Chilean coast in the Southern zone: Niebla (39.87°S 73.40°W, Los Lagos Region) and Coliumo (36.55°S 72.95°W, Biobío Region). The total biomass (20-30 kg of fresh tissue) was transported at 4° C to the laboratory, and washed with fresh water, posteriorly with 1 µm filtered-seawater and cleaned before freezing.

### 2.3 Oleoresin analysis

#### 2.3.1 A.chilense oleoresin production

Harvested seaweed was frozen at −20°C for two weeks and then chopped into small pieces (3-5 mm in length) by using a knife for the production of oxylipins induced by algal tissue after mechanical damage (22). Chopped alga was lyophilized to completely dry the biomass, and dried powdered seaweed was obtained after grinding in a coffee grinding machine. Dichloromethane (DCM) was used as an organic solvent for lipid extraction of pulverized dry seaweed. In brief, 50 g of pulverized dry alga was placed in an Erlenmeyer flask with 150 mL of DCM, shaken for 30 min and filtered. The remaining sediment was extracted once again as indicated above. The whole filtered extract was evaporated, suspended in cyclohexane and lyophilized to finally obtain a lipid extraction or oleoresin of *A. chilense (Gracilex®).* A detailed protocol is already published in the patent application WO/2014/186913 (61).

#### 2.3.2 Determination of total antioxidant capacity of the A. chilense oleoresin using a CUPRAC assay

Oleoresins obtained from dichoromethane extraction of lyophilized Spirulina (AquaSolar™) and Maqui (Isla Natura™) were prepared as described above for *Agarophyton chilense* extract production. The oleoresins were resuspended in DMSO, and 50 µLof sample was taken to measure the antioxidant capacity using an OxySelect™ TAC Assay kit (using the manufacturer’s instructions) (Cell Biolabs, INC. CA, USA), which is based on the reduction of copper(II) to copper(I) (62). Uric acid was used as a standard to calculate the milli-equivalents of antioxidant.

#### 2.3.3 Analysis of lipid and antioxidant contents of A.chilense oleoresin

Six different preparations of *Gracilex®* were used to measure tocopherols (α-tocopherol, γ-tocopherol and δ-tocopherol) and ß-carotene by HPLC-electrochemical methods. The fatty acid profile of *Gracilex®* was measured as fatty acid methyl esters by GC-FID. Both analyses were performed at the Center of Molecular Nutrition and Chronic Disease of Pontificial Universidad Catolica according to the Chilean normative procedure NCh2550-ISO5508.

### 2.4 Cellular studies

#### 2.4.1 Cell lines

HeLa, PC12 and 3T3-L1 cell lines were obtained from the American Type Culture Collection (ATCC; Manassas, USA). All the cells were cultured following ATTC recommendation in a 5% CO2 incubator at 37°C and 95% humidity.

PC12 cells were grown and cultured in RPMI 1640 medium supplemented with 10% horse serum (HS), 5% fetal bovine serum (FBS) and antibiotic/antimycotic (Ab-Am). HeLa cells were grown and cultured in DMEM high glucose with 10% FBS and penicillin/streptomycin, and 3T3-L1 cells were grown and cultured in DMEM high glucose with 10% fetal calf serum (FCS) and P/S. These cells were grown under low passages and at a maximum of 70% confluence.

#### 2.4.2 Cellular transfection

A PPAR reporter activity assay was performed by transient transfections of PC12 and HeLa cells with a reporter plasmid containing three tandem repeats of the peroxisomal proliferator response element (PPRE) fused to the herpesvirus thymidine kinase promoter upstream of the coding sequence for luciferase (PPRE-tk-LUC) and a pCMX vector containing full-length murine PPARγ1 (63). Cells were grown until 60–70% confluence in 24-well collagen-coated plates and transfected with 2 μL of Lipofectamine^TM^ 2000 (Thermo Fisher, MA, USA) and 0.3 μg of PPRE-tk-LUC vector, 0.26 μg of pCMX-PPARγ vector and 0.1 μg of β-galactosidase expression plasmid (CMV-β-Gal) (Clontech, now Takara Bio USA. Inc) vector for normalization.

HeLa cells and PC12 cells were seeded in 48-well plates at a density of 1×10^5^ cells/well in 250 µL of RPMI 1640 medium (10% HS, 5% SFB and Ab-Am) grown for 24 h and transfected with Lipofectamine 2000 (Thermo Fisher, MA, USA) with 0.8 µg of the reporter plasmid (Gal4-dependent MH100tk-Luc), 0.8 µg of the fusion protein plasmid (PPARγGAL4) and 0.053 µg of the CMV-β-Gal as an internal control (64, 65). Transfection was carried out in low serum RPMI 1640 medium (2% SFB, 2% HS) without antibiotics for 6 h, the medium was replaced, and the cells were treated with the indicated extracts and corresponding vehicle for 16 h. The medium was then discarded, and the cells were lysed with reporter lysis buffer (Cell Culture Lysis Buffer, Promega, WI, USA). The lysate was centrifuged, and the luciferase activity of the supernatant was determined using a luciferase assay system (Promega, MA, USA) and measured in a luminometer (Turner Biosystem 20/20). For normalization of luciferase activity, β-Gal activity was evaluated using o-nitrophenyl-β-D-galactopyranoside (ONPG) as the substrate in a standard colorimetric enzymatic assay. The luminescence signals obtained from the luciferase activity measurements were normalized to β-Gal colorimetric measurements to account for differences in the transfection efficiency or cell number. All transfection experiments were performed in triplicate.

#### 2.4.3 Adipocyte differentiation

3T3-L1 cells were differentiated as described previously (66). Briefly, 3T3-L1 cells were seeded in DMEM with 10% FCS at a density of 6 × 10^5^ cells/35 mm plate for lipid staining or 2× 10^5^ cells/12-well plate for RNA extraction. After 48 h, the differentiation medium was used; it contained freshly prepared solutions of 1 μg/mL insulin, 0.5 mM isobutylmethylxanthine (IBMX) and 0.1 µg/mL dexamethasone in the presence or absence of test compounds: rosiglitazone, RGZ (1 µM), FMOC-Leu (25 µM) or *A. chilense* oleoresin (60 µg/mL). After 48 h, the differentiation medium was replaced with DMEM containing 1 μg/mL insulin, and then, the medium was changed to DMEM supplemented with 10% FBS and P/S. Every 2 days the medium was changed. After 7 days of differentiation, the cells were harvested for RNA extraction using a HiBind column (Omega Bio-Tek, GA, USA) following the manufacturer’s instructions. On the 10th day of differentiation, the cells were stained with oil red O, as described previously. Briefly, the cells were washed twice with PBS at 37°C and then fixed with 4% paraformaldehyde (PFA) for 1 h at room temperature. The plate was washed with PBS and treated with 60% isopropyl alcohol for 6 min. After the plates were dried, oil red O (6:4) was added for 2 h at room temperature. Subsequently, the plates were washed extensively, dried and photographed. For quantification of oil red O incorporation in cells, the cells fixed in the plates were solubilized with pure isopropyl alcohol, and the absorbance of the solution was measured at 490 nm (67).

#### 2.4.4 MTT viability assay

The MTT (3-(4,5-dimethylthiazol-2-yl)-2,5-diphenyl tetrazolium bromide) assay was performed in PC12 cells as described before with minor modifications (68). Briefly, PC12 cells were seeded in 48-well plates at a density of 1.5×10^5^ cells/well in 250 µL of RPMI 1640 medium (10% HS, 5% SFB and Ab-Am) and grown for 24 h. Then, the medium was replaced with low serum RPMI 1640 medium (2% SFB, 2% HS, Ab-Am).

3T3-L1 cells were seeded in 48-well plates at a density of 9×10^4^ cells/well in 250 µL of DMEM (10% CS and Ab-Am) and grown for 24 h. The cells were treated with the indicated extracts and the corresponding vehicle for 16 h. The cells were washed with PBS and then incubated with 0.5 mg/mL MTT dissolved in medium without phenol red or serum for 2 h at 37°C. The MTT solution was discarded, and the cells were lysed with pure DMSO. The formazan product was measured in a spectrophotometer at 570 nm and 620 nm. The difference between both measurements was used to calculate the percentage of cell survival with respect to the control.

#### 2.4.5 RT-qPCR

RNA quality was evaluated by agarose gel integrity. Any contaminant genomic DNA was degraded by DNase treatment (RQ1, RNase-free DNase, Promega Corp., WI, USA). cDNA was generated using M-MLV reverse transcriptase (Promega Corp., WI. USA) and random primers. qPCR was performed using SYBR Green (Brilliant II SYBR Green qPCR Mastermix, Agilent Tech. CA, USA) in a Stratagene MX-3000P-qPCR System (Agilent Technologies, CA, USA). *Fabp4* (fatty acid binding 4) and *Lpl* (lipoprotein lipase) genes were amplified together with three reference genes in each run: glyceraldehyde-3-phosphate dehydrogenase (GAPDH), NoNo and β-actin. The murine primer sequences used for Fabp4 were F: 5’-GCGTGGATTTCGATGAAATCA-3’, R: 5’-CCCGCCATCTAGGGTTATGA-3’; for Lpl: F: 5’-ATTTGCCCTAAGGACCCCTG-3’, R: 5’-GCACCCAACTCTCATACATTCC-3’; for GAPDH: F: 5’-TGCACCACCAACTGCTTAGC-3’, R: 5’-GGATGCAGGGATGATGTTCT-3’; for NoNo: F: 5’-TGCTCCTGTGCCACCTGGTACTC-3’, R: 5’-CCGGAGCTGGACGGTTGAATGC-3’ and for β-actin: F: 5’-CCTGTGCTGCTCACCGAGGC-3’, R: 5’-GACCCCGTCTCCGGAGTCCATC-3’.

### 2.5 Mouse studies

#### 2.5.1 Mouse treatments

Fifty adult (6 weeks old) C57BL/6J mice were imported from the Jackson Laboratory (USA). The mice were housed in a room with controlled temperature (24°C) and humidity (55%-60%) for a 12 h light/dark regime. Water and food were supplied ad libitum by Pontificia Universidad Católica de Chile Catholic University Animal Facility. All of the animal procedures were conducted in accordance with the guidelines of the Chilean National Council for Science and Technology (CONICYT).

At 6 weeks of age, the mice were separated into 5 groups of 10 animals and fed two different diets. Four groups were fed a high fat diet (HFD: 60% kcal based on fat, Research Diet, Inc., USA), and one group was fed a low-fat diet (LFD: 10% kcal based on fat) (69, 70).

The animals were weighed twice a week throughout the study. At 12 weeks of age, the mice were treated as follows: the control diet (LFD) mice were treated with corn oil; the control high-fat diet (HFD) mice were treated with corn oil; the positive control HFD mice were treated with rosiglitazone (RZG) (5 mg/kg); the HFD mice were treated with two different doses of *Gracilex®*, 90 mg/kg and 300 mg/kg. The treatment was administered orally once a day by gavage for 30 days. The daily volume administered was approximately 50 µL.

#### 2.5.2 Measurement of plasma metabolic parameters

Blood samples were obtained at the beginning (12 weeks old) and at the end of the treatment (16 weeks old). Before sampling, the mice were fasted for 15 h. The sample was obtained by puncture of the submandibular sinus, collected in heparinized tubes, and immediately centrifuged (3500 rpm/10 min/4°C) to obtain blood plasma.

Mouse adiponectin (DuoSet® R&D Systems, USA) and mouse insulin (Ultrasensitive EIA, ALPCO Diagnostic, USA) were detected by ELISAs. Plasma levels of glucose and cholesterol were determined by using enzymatic chemical kits (Wiener Laboratories SAIC, Argentina).

#### 2.5.3 Histopatholgical studies

Twelve C57BL/6J mice (12 weeks old) per experimental group were randomly assigned to one of the following treatments: corn oil (control) or *Gracilex®* (300 mg/kg). The treatment was administered orally once a day by gavage for 15 days. The daily volume administered was approximately 50 µL. At the end of the treatment, the mice were euthanized by isoflurane overdose. The liver, kidney, esophagus and stomach were extracted and maintained in 9.25% formalin (4°C) for 48 h. For the histopathological evaluation, the samples were embedded in paraffin, processed under standard conditions and stained with hematoxylin-eosin (H/E). Liver fat accumulation was determined by Sudan black staining, and liver glycogen deposition and kidney basal membrane evaluation were determined by Schiff’s periodic acid (SPA) staining. The histopathological findings were classified according to magnitude (0 = absent; I = slight; II = moderate; III= severe).

### 2.6 C. elegans studies

Nematodes were raised at 20°C under standard laboratory conditions on agar plates cultured with Escherichia coli (OP50) as a food source (71). The oxidative stress-sensitive *msra-1(tm1421)* mutant strain (72) was use for stress resistant assays. Previously, for the oxidative stress challenge, the nematodes were fed for 24 h with *Gracilex®* or its vehicle DMSO. For administration of *Gracilex®* to the nematodes, plates were prepared as follows: *Gracilex®* was dissolved in DMSO at a concentration of 100 mg/mL. A stock of dead bacteria was prepared by growing the *E. coli* OP50 strain in LB buffer overnight. After centrifugation, the pellet was resuspended to obtain a stock concentrated 6 times. This stock was heated at 70°C for 30 min. The stock of dead bacteria was stored at −20°C until use. Twenty-five micrograms of *Gracilex®* were mixed with 50 µL of dead bacteria and NP40 (to a final concentration of 0.1%) to a final volume of 52 µL. This mix was seeded on 2 mL of NGM agar in a 35 mm plate. The plates were left to dry in the dark ON in the laminar flow chamber and stored at 4°C in the dark for no more than 1 week before use. For the oxidative stress test, L4 stage *msra-1* nematodes were growth for 24 h on plates with *Gracilex®* or DMSO (vehicle). On the day of the test, the nematodes were transferred to a 24-well plate containing 600 µL of 9 mM hydrogen peroxide dissolved in M9 medium (minimal medium containing minimal salts: KH2PO4, 15 g/L, NaCl, 2.5 g/L, Na2HPO4, 33.9 g/L, NH4Cl, 5 g/L). Ten nematodes were placed per well, and the movement was monitored every 10 min as described in (73) until there was no living nematode (approx. 2 h). Each experiment was performed in triplicate (30 nematodes per condition) (74).

### 2.7 Statistical analysis

Cell-based assays for PPARγ transactivation were performed in triplicate and in at least three independent experiments. The results are expressed as the mean fold induction relative to the control condition ±SEM. One-way analysis of variance (ANOVA) followed by Tukey’s multiple comparison tests was applied to determine statistical significance between the control and treatment conditions. The same statistical tests were used for the analysis of 3T3 adipocyte differentiation in the oil ed O assay and in lipogenic gene mRNA induction determined by qPCR. For *in vivo* studies of insulin resistance, an unpaired t test was used to compare the low-fat diet group and the HFD group. For analysis of the differences between subgroups of HFD-fed mice and LFD-fed mice, one-way ANOVA and Tukey’s multiple comparison tests were applied. Finally, in *C. elegans in vivo* experiments, two-way ANOVA with repeated measures was used to compare differences in the survival curves of the nematodes fed *Gracilex®* and the nematodes under control conditions. Fisher’s LSD post-test was used to determine significant differences at appropriate times. Significant differences were calculated and are represented with the following symbols of significance level: * *p* < 0.5, ** *p* < 0.01, *** *p* < 0.001. Statistical analysis was performed using GraphPad Prism 5.0 for Mac (GraphPad Software, San Diego, CA, USA).

## 3 Results

### 3.1 Effect of A. chilense oleoresin on PPARγ transcriptional activity

The oleoresin or lipid extract derived from *A. chilense* (*Gracilex®*) was produced as described in patent application WO/2014/186913 (61) (see methods for further details). To determine whether *Gracilex®* can induce the transcriptional activation of PPARγ, we used two different cell lines, HeLa and PC12, for transient transfection assays using the PPRE-TK-LUC reporter (75) cotransfected with the PPARγ expression vector (Fig. 1A). In both cell types, the *Gracilex®* induced significant activity at 40 μg/mL, resulting in an activation of up to 65% of the maximal activation achieved by the full agonist RGZ at 1 μM. To further confirm the activation of PPARγ by lipids derived from *A.Chilense* and to study the contribution of PPARγ to the increased luciferase, we assessed the ability of *Gracilex®* to activate the chimeric GAL4-DBD-PPAR-LBD protein from a Gal4-dependent MH100-Luc reporter using PC12 cells (Fig. 1B). In this assay, the activation of the reporter is activated by the chimeric GAL4-DBD fusion protein, thus avoiding potential interference from any endogenous receptor (64, 65). We observed dose-response activation of PPARγ with *Gracilex®*. At a concentration of 100 μg/mL, *Gracilex®* showed PPARγ activity induction comparable to that promoted by the SPARMs INT131 and FMOC-Leu at 25 μM (Fig. 1B). In contrast, RGZ-induced PPARγ activation was greater than 8 times higher at 1 μM than that of *Gracilex®*, which is consistent with the definition of a full agonist (Fig. 1B). The PPARγ activation induced by *Gracilex®* was prevented upon cotreatment with the PPARγ antagonist T0070907 (76), demonstrating that *A. chilense* lipids specifically activate PPARγ (Fig. 1C). These results suggest that *Gracilex®* is a good source of natural ligands for PPARγ. No effects on the survival of PC12 cells were observed for any of the *Gracilex®* concentrations evaluated (Supplementary Fig. 1A). SPPARMs are selective modulators of PPARγ activity with partial agonism and reduced side effects such as adipose tissue gain (77). Considering that PPARγ is a master gene for adipocyte differentiation and *Gracilex®* activates PPARγ similarly to SPPARM at the higher doses evaluated (60 μg/mL and 100 μg/mL), we next characterized the effect of *Gracilex®* on 3T3-L1 cellśs adipocyte differentiation using a well-established protocol (66). The incubation of the 3T3-L1 cell line with RGZ (1 μM) and a hormonal mix for seven days induced an evident increase in oil red O staining, which labels lipid droplets, indicating the accumulation of triglycerides (78). The quantification by absorbance at 490 nm of the isopropanol soluble fraction (67) derived from 3T3-L1 differentiated cells showed a twofold increase in the cells treated with RGZ compared to the cells treated with only a hormonal mixture (Fig. 2A, B). In contrast, neither FMOC-Leu nor INT131 at 25 μM or *Gracilex®* at 60 μg/mL mimicked the effects of RGZ (Fig. 2A, B). To further confirm that SPPARMs and *Gracilex®* do not induce adipose differentiation, we measured the induction of mRNA levels of the adipogenic marker genes *lipoprotein lipase* (*LPL*) and *fatty acid binding protein 4* (*FABP4*). We observed that only RGZ promoted the adipocyte differentiation program in 3T3-L1 cells (Fig. 2C). To uncover the possible toxic effects of *Gracilex®* in differentiated 3T3-L1 cells, we determined cell viability by the MTT assay after *Gracilex®*, FMOC-Leu, INT131 or RGZ treatment. No changes in cell survival were found for each PPARγ agonist in 3T3-L1 cells (Supplementary Fig. 1B). Taken together, these results indicate that *Gracilex®* can induce PPARγ activation in a similar fashion to SPPARMs without promoting adipocytic differentiation or toxic effects in 3T3-L1 cells at the concentration used.

**Figure 1.**
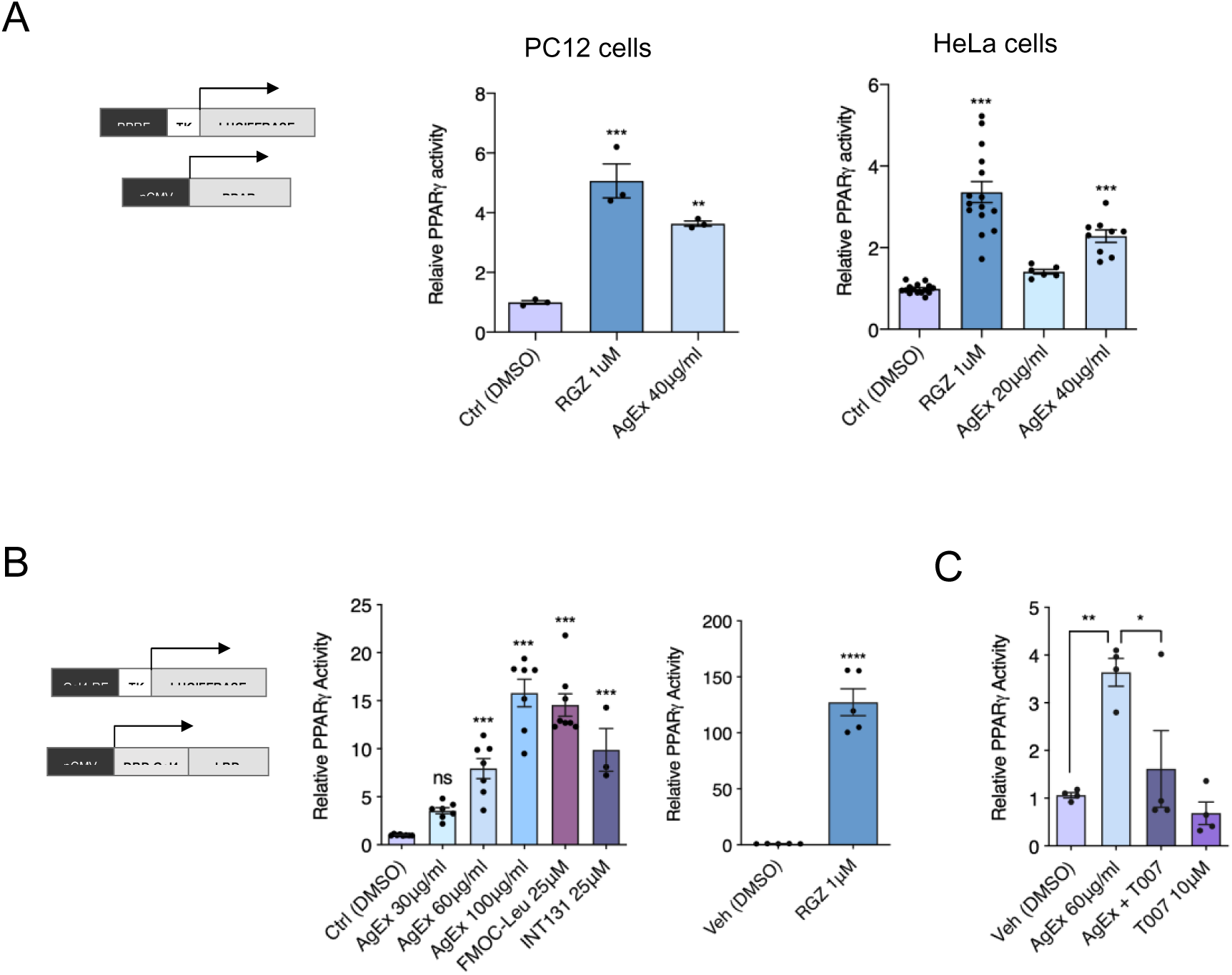
*A. chilense* oleoresin induces PPARγ transcriptional activation. Two different cell-based reporter assays to measure PPARγ activation were performed. (**A**) Left panel, schematic representation of vectors used for cotransfection. PC12 and HELA cells were cotransfected with plasmids expressing full-length murine PPARγ and a reporter plasmid containing PPRE elements driving the expression of luciferase, together with a plasmid expressing beta-galactosidase as a transfection control. The full PPARγ agonist rosiglitazone (RGZ, 1 µM) and the *A.chilense* oleoresin or *Gracilex® (AgEx*, 40 µg/mL) induce the transcriptional activation of PPARγ, as shown by the increased luciferase activity and normalized to beta-galactosidase expression (expressed as relative PPARγ activity). In PC12 cells (n=3) and HeLa cells (n=6-13), the treatments (RGZ and *AgEx*) were compared to control transfected cells (DMSO), and the values are presented as ±SEM, ** *p*<0.01 \*\*\**p*<0.001 (one-way ANOVA, Tukey’s post hoc test). (**B**) Left panel, schematic representation of vectors used for cotransfection. A plasmid with expression of a chimeric construct of the PPARγ ligand binding domain (PPARγ LBD) gene and the DNA binding domain of Gal4 (Gal4LBD) was cotransfected with the response element of GAL4 (REGal4) driving the expression of luciferase. As in A, a third plasmid was included with expression of beta-galactosidase as a transfection control. Increased PPARγ activation is expressed as relative PPARγ activity. *AgEx* induces a dose-response activation of PPARγ comparable to the activation induced by the selective SPPARMs of PPARγ, FMOC-LEU and INT131 at a concentration of 25 µM. Instead, RGZ at 1 µM shows much higher activation (110 times over the control), as expected for a full agonist. Values are presented as ±SEM, n=3-7, \*\*\**p*<0.001 (left panel, one-way ANOVA, Tukey’s post hoc test. Right panel, unpaired t test). (**C**) PC12 cells were transfected as described in B and treated with *AgEx* (60 µg/mL) with or with the PPARγ antagonist T0070907 (T007, 10 µM). Values are presented as ±SEM, n=4, \**p*<0.5 (one-way ANOVA, Tukey’s post hoc test).

**Figure 2.**
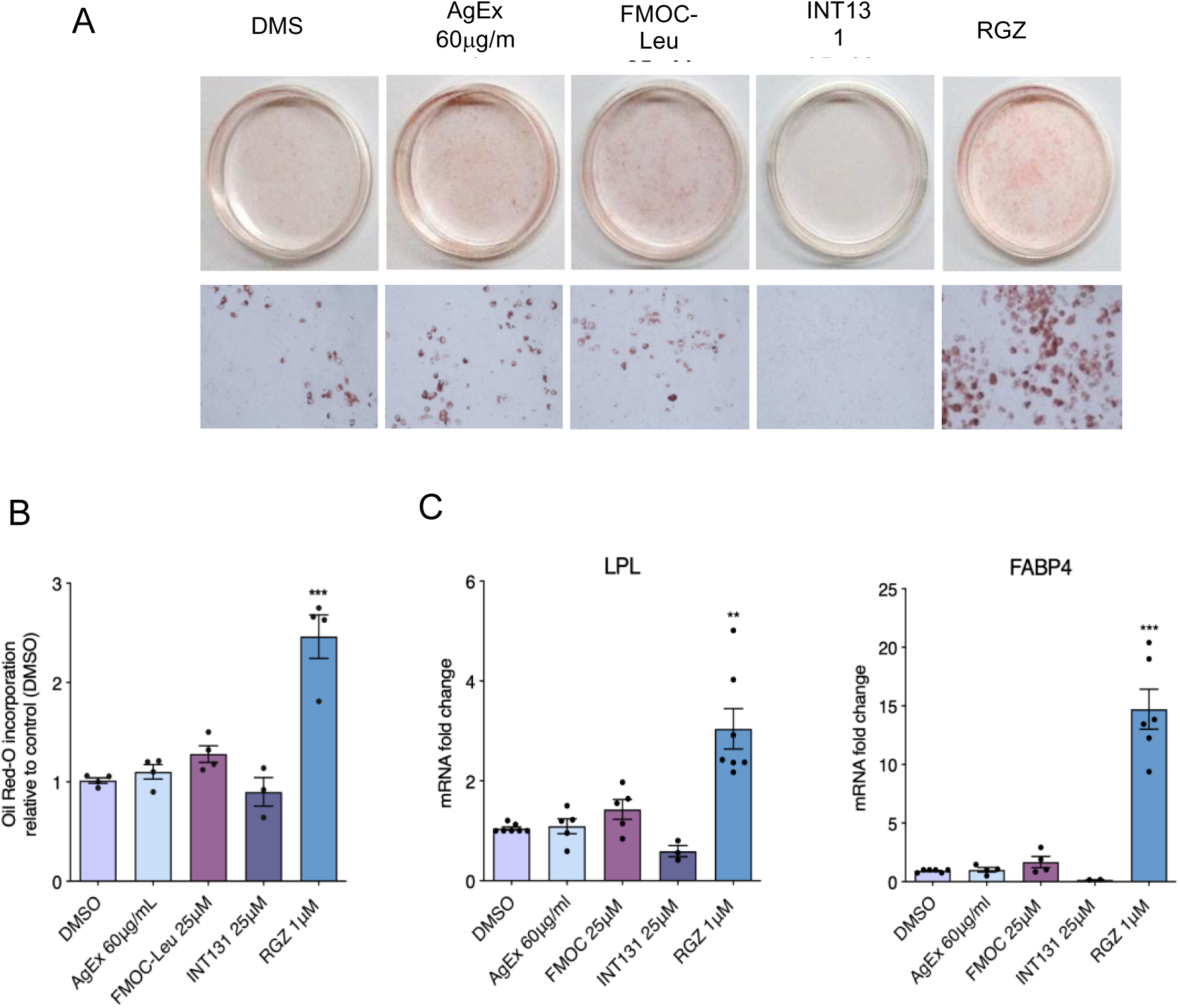
Adipogenesis in 3T3-L1 fibroblast cells is induced by rosiglitazone but not by *A. chilense oleoresin* or by the PPARγ SPPARMs FMOC-Leu and INT131. (**A**) First, 3T3-L1 preadipocytes were cultured with or without each PPARγ agonist, FMOC-Leu and INT131 (25 μM); RGZ (1 μM) and *A. chilense oleoresin* or *Gracilex®* (*AgEx) at* 60 μg/mL as specified in the methodology section. After 10 days of culture initiation, the cells were stained with oil red O and triglycerides, and the plates were photographed. (**B**) Quantification of oil red O incorporation was measured after cell lysis with isopropanol at a wavelength of 490 nm. Values are expressed as the mean relative to that of the control cells (treated with DMSO) ±SEM, n=6, \*\*\**p*<0.001 (one-way ANOVA, Tukey’s post hoc test). (**C**) Effects of *AgEx* and PPARγ agonists on the expression of lipogenic pathway-related genes during the differentiation process of mouse 3T3-L1 preadipocytes were measured after 7 days of culture initiation as indicated in the methodology section. After treatment with each compound, the cells were lysed, and the mRNA levels of the *LPL* and *FABP4* genes were measured by qPCR. The graph shows the relative abundance compared to the control (untreated cells, control DMSO). Values are expressed as the mean relative to that of the control cells (treated with DMSO) ± SEM, n=4, ∗∗*p*<0.01, ∗∗∗*p*<0.001 (one-way ANOVA, Tukey’s post hoc test).

### 3.2 Effect of A.chilense oleoresin on the metabolic dysfunction caused by high-fat diet (HFD)-induced obesity in male mice

Although there are lean individuals with T2DM, obesity explains 60-80% of diabetes cases (52–54). Even though animals do not develop T2DM like humans, there are appropriate animal models for studying the effects of new natural products such as *Gracilex®* on the development of T2DM induced by a high-fat diet. Studies indicate that a diet high in saturated fat (and not, for example, a diet rich in sucrose) is the most appropriate for the development of metabolic dysfunction in murine models (79), imitating the physiological conditions associated with T2DM in humans, such as insulin resistance due to failure of pancreatic cells (59). The C57BL/6J strain is a very useful strain for the development of diabetes induced by a high-fat diet (HFD) compared to other strains because it is genetically predisposed to develop obesity (69, 80). This strain shows two critical characteristics of T2DM, insulin resistance and dysfunction of pancreatic beta cells. These animals are an ideal model for studying new therapeutic agents, including natural products, and their potential to modulate this condition (59). Therefore, we used this model to test the potential use of *Gracilex®* as a nutraceutical agent to improve insulin and glucose resistance induced by an HFD. Initially, we compared the body weight gain between a group of C57BL/6J mice fed an HFD and another group fed a normal diet (low fat diet, LFD). After 12 weeks, a significant difference (*p* <0.001) in body weight was observed between the HFD and LFD groups, being 27% greater in the high caloric diet group than in the group receiving a normal diet (Fig. 3A). When we compared biochemical blood parameters in LFD vs HFD mice, a significant increase in glucose (81.7 v/s 119.5 mg/dL) (Fig. 3B) and insulin (0.29 v/s 0.62 ng/mL) (Fig. 3C) levels was observed, consistent with the insulin metabolic dysfunction. Additionally, the values of plasma adiponectin were markedly decreased (3.8 vs 1.8 μg/mL) in the HFD mice compared to the LFD mice (Fig. 3D), confirming the parameters described in obese animals with insulin resistance. Finally, as a result of high fat intake, plasma cholesterol values increased significantly (Fig. 3E) in the HFD vs LFD mice (121 v/s 152 mg/dL). Taken together, these results indicate the effective induction of the insulin resistance state by HFD intake in mice, which is the condition required to test the effect of *Gracilex®*.

**Figure 3.**
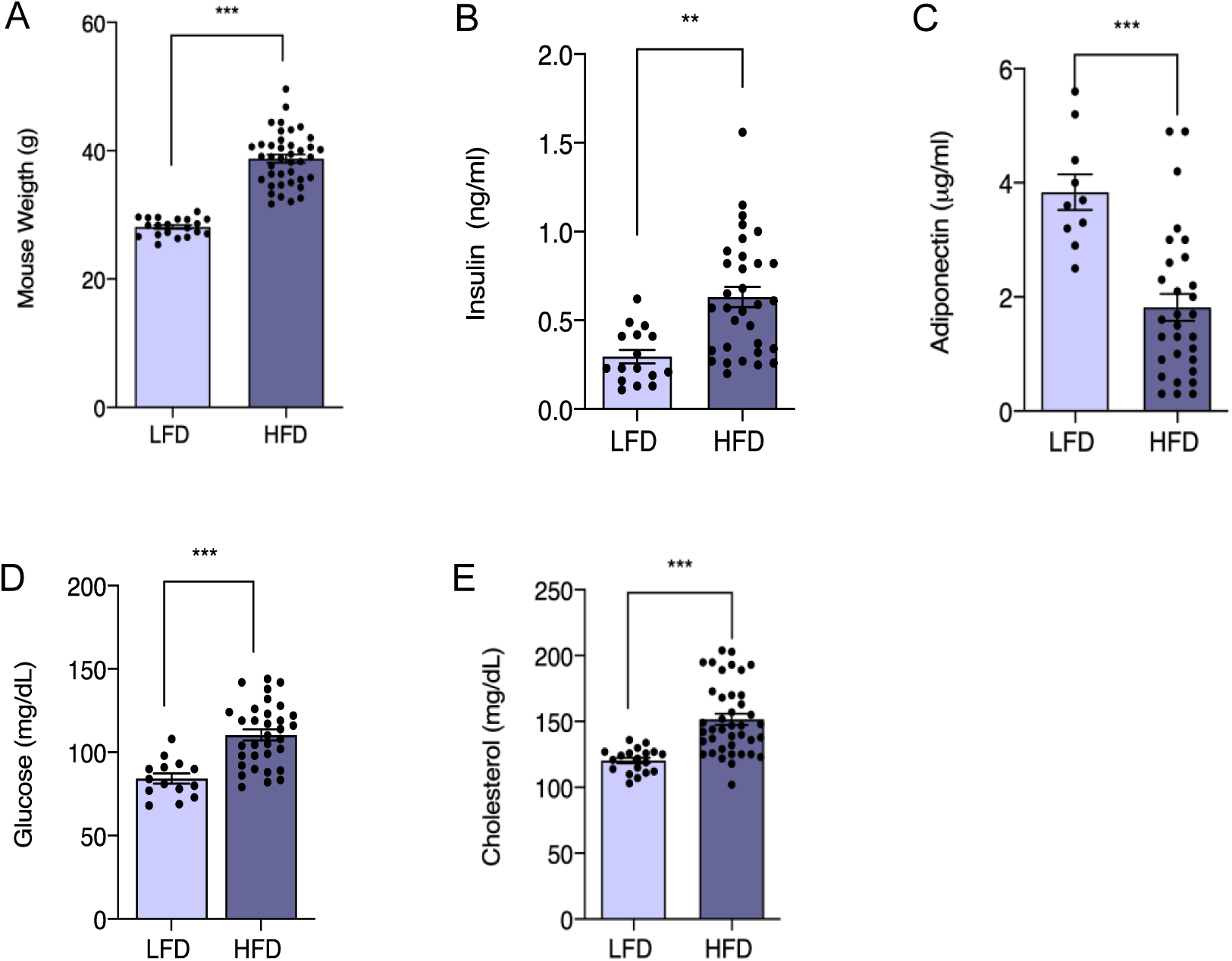
Promotion of weight gain and glucose and insulin resistance in male C57BL/6J mice administered a high-fat diet (HFD). Six-week-old mice were treated for an additional 6 weeks with a low-fat diet (LFD, n=20) and HFD (n=40). (**A**) Weight recording in groups of mice given an LFD and HFD after six weeks of treatment. During the whole period, the weights were recorded twice a week. Values are expressed as the mean recorded weight ± SEM, \*\*\**p* <0.001 (unpaired t test). (**B-E**) Biochemical blood parameters were measured in both groups: plasma glucose (mg/dL) (B), plasma insulin (ng/mL) (C), plasma adiponectin (μg/mL) D, and plasma cholesterol (mg/dL) E. Values are expressed as the mean± SEM, and significant differences are highlighted between the LFD vs HFD group ** *p*<0.01, \*\*\**p*<0.001 (unpaired t test).

To evaluate whether possible harmful effects of *Gracilex®* ingestion may be observed in mice, we performed a histopathological study in mice treated for a shorter period of time (15 days). No major changes in the kidney and stomach were found (data not shown). In the liver, we observed moderate glycogen accumulation by PAS staining, and no changes in fat accumulation in the mice fed a normal chow diet (Supplementary Fig. 2A-D). Additionally, no changes in body weight were found (Supplementary Fig. 2E). These results suggest no toxic effects were induced by *A. chilense* oleoresin.

The HFD-treated mice were then subdivided into four groups, maintained in an HFD and orally treated for 30 days with RGZ at 5 mg/kg/day and two different *Gracilex®* doses (90 mg/kg/day, and 300 mg/kg/day) or corn oil as vehicle. After the treatment period, no variations in body weight were observed between the subgroups; however, differences between the HFD group and the LFD group were maintained (Fig. 4A), as demonstrated previously (69). The analysis of blood parameters at the end of the treatment period showed that the mice receiving the HFD and corn oil maintained a significant increase in glycemia compared with the controls (Fig. 4B). In the HFD-fed mice, the group treated with *Gracilex®* (300 mg/kg) showed a complete recovery of blood glucose levels, reaching the levels of the LFD control mice (Fig. 4B). RGZ and *Gracilex®* (90 mg/kg and 300 mg/kg) were also able to decrease glucose to control levels (Fig. 4B). In relation to insulin parameters, the HFD mice treated with *Gracilex®* at a dose of 300 mg/kg showed a significant decrease in insulin, reaching values close to 50% of the concentrations found in the insulin-resistant HFD-treated control group (Fig. 4C). However, as expected (70, 81), synthetic RGZ promoted a decrease in insulin levels, normalizing this parameter to that observed in the control LFD mice

**Figure 4.**
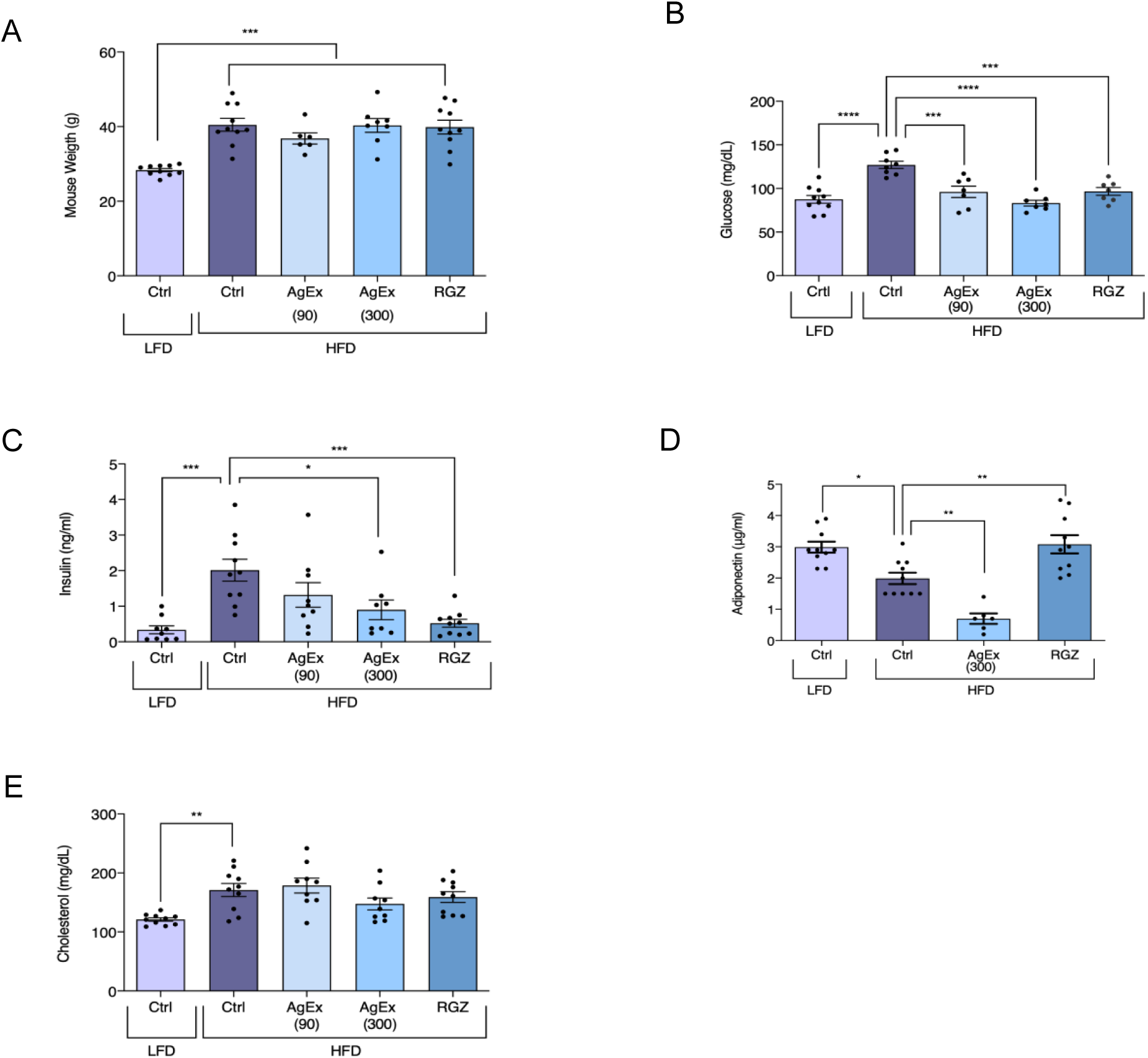
*A. chilense* oleoresin improves altered glucose and insulin parameters in the plasma of the male C57BL/6J mice given an HFD. After 6 weeks of LFD and HFD treatments as indicated in Fig. 3, LFD mice continued with the same diet with additional daily treatment with approximately 50 µL of corn oil (vehicle of drugs) for one month (30 days). The HFD-treated mice were divided into 4 groups, and they continued with the same diet, but 4 different daily treatments were given: corn oil (vehicle), 90 and 300 (mg/kg) of *A. chilense oleoresin* or *Gracilex®* (*AgEx)* and RGZ (5 mg/kg). During the whole period, the weights were recorded twice a week. (**A**) Comparison of weights at the beginning and at the end of the treatment for each group. Values are expressed as the mean ± SEM, n=10, \*\*\**p* <0.001 (one-way ANOVA, Tukey’s post hoc test). (**B-E)**, After the treatment, the mice were fasted for 15 h before taking blood samples, and plasma glucose (mg/dL) (**B**), plasma insulin (ng/mL) (**C**), plasma adiponectin (μg/mL) (**D**), and plasma cholesterol (**E**) (mg/dL) are shown. Values are expressed as the mean ± SEM, n=10. Significant differences are highlighted between the LFD and HFD groups (left panels** *p* <0.01, *** *p* <0.001, unpaired t test). Significant differences found in post-treatment experiments in the HFD group are regarding their HFD control. * *p* ≤0.05; ** *p* <0.01 *** *p* <0.001 (one-way ANOVA, Tukey’s post hoc test).

Adiponectin levels completely recovered after 30 days of RGZ treatment in the HFD-control mice compared with the HFD-RZG mice, which was equivalent to the values measured in the LFD control group (Fig. 4D). Unexpectedly, *Gracilex®* (300 mg/kg/day) had an opposite effect compared to RGZ, and a significant reduction in adiponectin levels was found compared to that of the HFD-control mice and the HFD-*Gracilex®* (300 mg/kg) mice (Fig. 4D), which is consistent with the fact that *Gracilex®* does not induce adipocyte differentiation in 3T3L1 cells like RZG (Fig. 2). Finally, plasma cholesterol was not affected either by treatment with RGZ or with *Gracilex®* in the HFD-treated mice (Fig. 4E), and this group maintained high levels compared to those of the group of mice fed a normal diet. These data demonstrated that *A.chilense* oleoresin improves insulin sensitivity and normalizes glucose tolerance with no alterations in weight gain, despite the reduced levels of circulating adiponectin, which is an adipose tissue-derived hormone that regulates the expansion and healthy function of adipose tissue (60). According to our studies, it is possible to consider *Gracilex®* as an alternative source of natural PPARγ ligands with similar properties found in selective PPAR modulators, presenting partial receptor activation that is sufficient for insulin sensitization in HFD-induced metabolic dysfunction with no effect on the promotion of adipocyte differentiation.

Species of the *Gracilaria* genus have already been studied in terms of lipid components by several authors, highlighting the rich and unique composition of sulfoglycolipids, polyunsaturated fatty acids, oxidized fatty acids, pigments and lipophilic antioxidant molecules (15,18,21,82,83). An overall analysis of some lipid components was performed in dichloromethane oleoresin prepared from *A. chilensis*. The *Gracilex®* fatty acid distribution is indicated in Table 1 as the relative abundance % of total fatty acid methylesters detected by gas chromatography with a flame ionization detector (GC-FID). We found a high content of saturated fatty acids (51.4 ± 4.89%), followed by polyunsaturated (25.7 ±3.12%) and monounsaturated (19.6 ±2.88%) fatty acids. The most abundant fatty acids were palmitic acid (C16:0), which contributed 40% of the total fatty acids analyzed; ARA (C20:4, n-6), 21.1%; oleic acid (C18:1, n-9), 14.13%; myristic acid (C15:0), 4.4%; and 4.0% of 11-octadecenoic acid (C18:1, n-7) (Table 1). In general, we found that this distribution is similar to the fatty acid profile reported by Da costa et al. (18) in their methanolic extract of *Gracilaria* sp., a red seaweed closely related to *A. chilensis*. The omega-3 fatty acid content was low, reaching up to 1.22% of the total fatty acids analyzed. In contrast, the total content of omega-6 fatty acids was higher (Table 1). ARA is the precursor of eicosanoids and oxylipins, bioactive lipidic promoters of the defense response in seaweed (19, 20). Most natural and endogenous PPARγ ligands are derived from ARA metabolization (84); therefore, the increased abundance in this seaweed extract may account for its high PPARγ transactivation capacity.

**Table 1.** Fatty acid composition (relative abundance, % over total fatty acids) of *A. chilense oleoresin*.

|  | Fatty Acid | Chain Lenght | Mean % | SD |
| --- | --- | --- | --- | --- |
| Saturated | Decanoic Acid | 10:00 | 0,760 | 1,1 |
|  | Dodecanoic Acid | 12:00 | 0,305 | 0,3 |
|  | Tridecanoic Acid | 13:00 | 0,983 | 0,3 |
|  | Tetradecanoic Acid | 14:00 | 4,438 | 0,9 |
|  | Pentadecanoic Acid | 15:00 | 0,440 | 0,3 |
|  | Hexadecanoic Acid | 16:00 | 40,005 | 5,4 |
|  | Heptadecanoic Acid | 17:00 | 0,750 | 1,3 |
|  | Octadecanoic Acid | 18:00 | 2,683 | 3,0 |
|  | Eicosanoic Acid | 20:00 | 0,152 | 0,08 |
|  | Docosanoic Acid | 22:00 | 0,238 | 0,05 |
|  | Tetracosanoic Acid | 24:00 | 0,127 | 0,08 |
| Mono-Insaturated | 10-Pentadecaenoic Acid | 15:1 n-5 | 1,87 | 1,8 |
|  | 9-Hexadecaenoic Acid | 16:1 n-7 | 0,46 | 0,3 |
|  | 9-Octadecaenoic Acid | 18:1 n-9 | 14,13 | 4,6 |
|  | 11-Octadecaenoic Acid | 18:1 n-7 | 4,07 | 1,4 |
| Omega-6 Polyunsaturated | 9,12-Octadecadienoic Acid | 18:2 n-6 | 2,87 | 0,81 |
|  | 6,9,12-Octadecatrienoic Acid | 18:3 n-6 | 0,17 | 0,10 |
|  | 11,14-Eicosadienoic Acid | 20:2 n-6 | 0,34 | 0,37 |
|  | 8,11,14-Eicosatrienoic Acid | 20:3 n-6 | 0,47 | 0,09 |
|  | 5,8,11,14-Eicosatetraenoic Acid | 20:4 n-6 | 21,06 | 3,81 |
| Omega-3 Polyunsaturated | 9,12,15-Octadecatrienoic Acid | 18:3 n-3 | 0,390 | 0,42 |
|  | 5,8,11,14,17-Eicosapentaenoic Acid | 20:5 n-3 | 0,408 | 0,38 |
|  | 7,10,13,16,19-Docosapentaenoic Acid | 22:5 n-3 | 0,27 | 0,12 |
|  | 4,7,10,13,16,19-Docosahexaenoic Acid | 22:6 n-3 | 0,125 | 0,04 |
| Other | Conjugated Fatty Acids |  |  |  |
|  | Fatty Acid | Chain Lenght | Mean % | SD |
|  | c9, t11-octadecadienoic Acid | 18:2 n-cla | 0,187 | 0,06 |
|  | Trans Fatty Acids |  |  |  |
|  | Fatty Acid | Chain Lenght | Mean % | SD |
|  | 10-Transpentadecaenoic Acid | 15:1 n-5t | 0,258 | 0,24 |
|  | 9-Octadecaenoic Acid | 18:1 n-9t | 1,037 | 1,30 |
|  | 11-TransOctadecaenoic Acid | 18:1 n-7t | 0,540 | 0,30 |
|  | 9,12-Octadecadienoic Acid | 18:2 n-6tt | 0,11 | 0,04 |

Fatty acids were detected as methyl fatty acid by GC-FID. Mean of six different lipid extracts produced from independent alga biomass which has been harvested and cultured in different seasons (fall, winter and summer).

The nutritional value of red macroalgae is also due to the diverse content of pigments and antioxidants (14,82,85). Consequently, we determined the total antioxidant capacity (TAC) using a CUPRAC assay (cupric ion reducing antioxidant capacity) that is compatible with lipophilic antioxidants (such as tocopherols) (62) in the oleoresin of *A. chilense* and in two different botanical lipid extracts, spirulin and maqui, produced with dicloromethane extraction as *Gracilex®*. Spirulin and maqui berries are known functional foods with high nutraceutical value. Spirulin is a cyanobacteria with known antioxidant and nutritional beneficial effects (86), and maqui berry (*Aristotelia chilensis*), a flavonoid-rich berry, is well known for its antioxidant and insulin sensitizing effects (87). Table 2 shows that *Gracilex®* is slightly superior to the other botanical oleoresins, reaching a mean of 430 mg uric acid equivalent per 100 mg of oleoresin. To further explore the antioxidant molecules, we evaluated the contents of tocopherols and ß-carotene in *Gracilex®*. As indicated in Table 3, we found high levels of ß-carotene and tocopherols, while lycopene was not detected. The beta-carotene content was approximately 1538 μg/g oleoresin, while the total tocopherol content was 6673 μg/g oleoresin, which was distributed as follows in the different types of tocopherols: α-tocopherol (527.7 μg/g), γ-tocopherol (5332.8 μg/g), and ∂-tocopherol (2660 μg/g) and. Thus, the antioxidant capacity of *Gracilex®* is partly due to the high levels of ß-carotene and tocopherols, particularly γ-tocopherol.

**Table 2.** Total antioxidant capacity (TAC) of *A. chilense oleoresin* and other botanical sources by CUPRAC.

| Sample | mg Uric Acid Eq /100mg Oily extract |  |  |
| --- | --- | --- | --- |
|  | Mean | SD | n |
| *A.chilense* oleoresin | 430 | 154,2 | 7 |
| Spirulina oleoresin | 344 | 156,9 | 3 |
| Maqui oleoresin | 305 | 86,4 | 3 |

**Table 3.** Antioxidant content characterization of *A.chilense* oleoresin.

| Antioxidant | µg/g of oleoresin |
| --- | --- |
|  | Mean ± S.E.M. |
| α-Tocopherol | 527.7 ± 85.3 |
| γ-Tocopherol | 5332.8 ± 1523.3 |
| δ-Tocopherol | 2660 ± 397.1 |
| Total Tocopherols | 6673 ± 1568.2 |
| β-Carotene | 1538 ± 378.4 |

Tocopherol and $\beta$ -carotene composition in *A.chilense oleoresin*. Value are given as the mean +/- S.E.M (n= 6 independent oleoresin preparations).

### 3.3 Antioxidant properties of A. chilense oleoresin

Considering the high contents of tocopherols and ß-carotene, we next assessed the ability of *Gracilex®* to confer antioxidant resistance *in vivo*. The oxidative stress-sensitive *msra-1* mutant strain of *C. elegans* (72) was chosen as a model to evaluate *in vivo* oxidative stress resistance. Since this strain has been proved to be very sensitive to acute oxidative treatment is a powerful tool to assess the physiological consequences of oxidative stress *in vivo* (88, 89). Taking advantage of the autofluorescence of pigments contained in *Gracilex®*, we first evaluated whether the nematodes were able to ingest *Gracilex®* in a mixture with dead OP50 bacteria. As shown in Fig. 5A, *Gracilex®* was visualized along the digestive track of the nematodes (highlighted by red arrows), indicating that they can ingest the *A. chilense* oleoresin. For antioxidant resistance assays, msra-1 L4 larval stage nematodes were fed for 24 h with DMSO (CTL), *Gracilex®* or N-acetylcysteine (NAC) mixed with dead bacteria. Then, the worms were challenged with a hydrogen peroxide solution, and their movement was scored every 10 min. Fig. 5B shows the percentage of active nematodes over time after the challenge. After 80 min, 60% of the *msra-1* mutants were already inactive compared to 40% of the nematodes fed *Gracilex®*. From 70-110 min, we consistently found more active nematodes, as shown by the right shift in the survival slope, which was significant at 80 min. Similar behavior was observed in the worms that were previously expose to NAC, a potent well-known antioxidant, at doses of 5 mM (Fig. 5C). These findings indicated that nematodes fed with *Gracilex®* extract presented increased resistance against oxidative stress, improving their survival in agreement with the CUPRAC analysis and the characterization of their antioxidant components.

**Figure 5.**
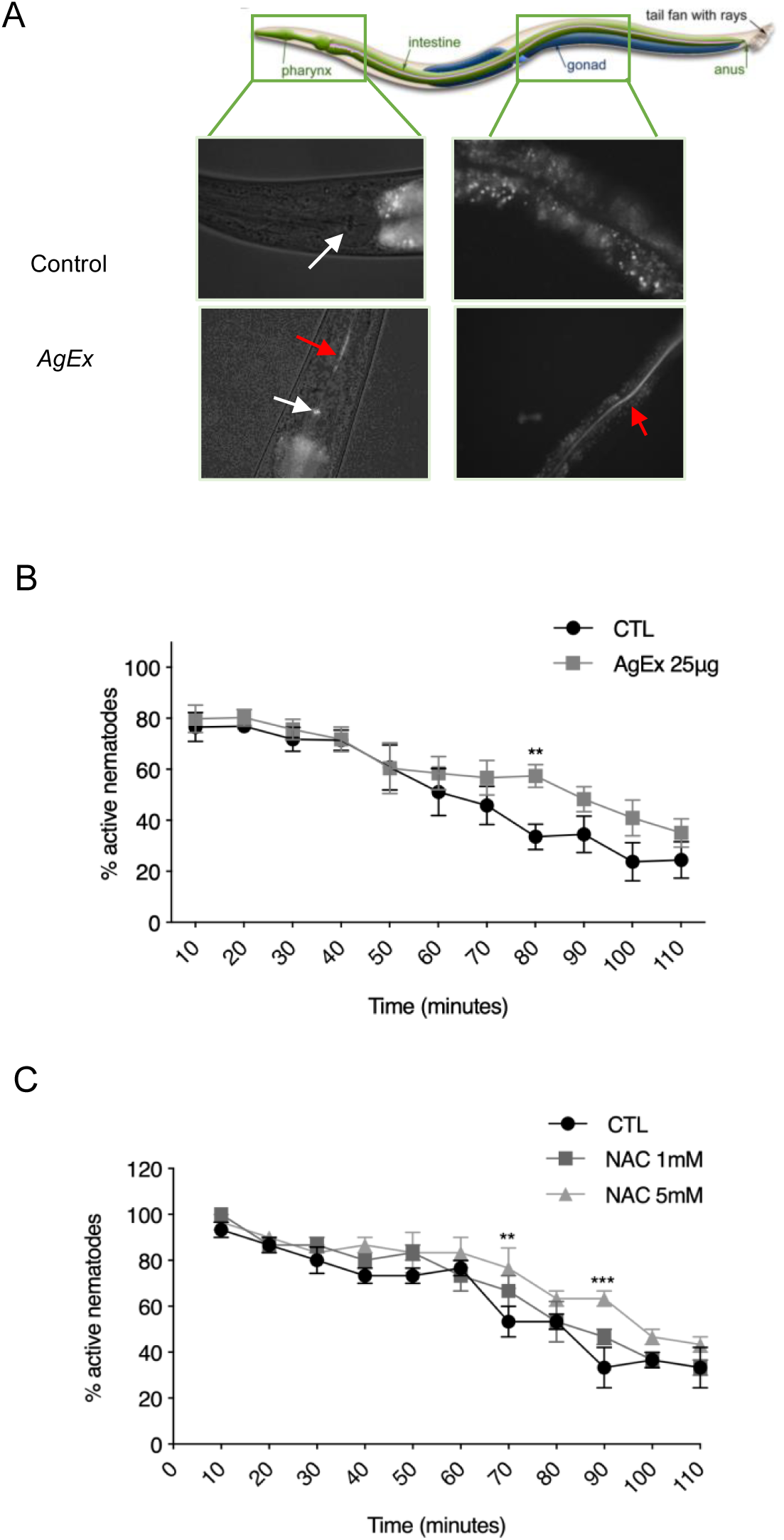
*A. chilense* oleoresin increases oxidative stress resistance in *C. elegans* challenged with hydrogen peroxide. (**A**) Representative illustration of a whole *C. elegans* organism extracted from Corsi A. K and cols., 2015 (125). Red autofluorescence of the extract was observed along the nematode digestive track visualized under fluorescence microscopy. *A. chilense oleoresin* or *Gracilex®* (*AgEx*) was dissolved in DMSO, mixed with 25 µL of dead bacteria and dispersed on 1.5 mL of agar. The white arrow shows the nematode pharynx. The red arrow highlights the fluorescent extract inside the nematode. Control corresponds to an adult N2 worm grown under standard conditions. (**B**) L4 stage nematodes from the msra-1 strain were fed *AgEx* for 24 h and then exposed to oxidative stress by 9 mM hydrogen peroxide for up to 110 min. Values are expressed as the mean (n=5) percentage of survival ± SEM, \*\**p*<0.01 (two-way ANOVA with repeated measures and Fisher’s LSD test). (**C**) The same experimental assay was performed as described in B, but nematodes were fed two different concentrations (1 mM and 5 mM) of N-acetylcysteine (NAC). NAC was used as a positive control. Values are expressed as the mean (n=3) percentage of survival ± SEM, \*\**p*<0.01 and \*\*\**p*<0.001 (two-way ANOVA with repeated measures and Fisher’s LSD test). Significant differences in the percentage of active nematodes were found under treatment with 5 mM NAC compared to the control at 70, 90 and 120 min after exposure to hydrogen peroxide.

## 4. Discussion

Interest in the study of natural bioactive molecules able to activate PPARγ has grown over the years due to the importance of the physiological processes that these receptors regulate and to the side effects caused by pharmacological PPARγ drugs. This interest is paralleled with an increase in the study of the biomedical potential of seaweed due to the large amounts of bioactive molecules they possess (11,90–92). The study of the lipid components of *A. chilense* and other closely related red seaweeds had demonstrated the presence of PUFAs and eicosanoids (19,22,37); however, whether the lipids present in the *Agarophyton* genus were able to activate PPARγ and had insulin sensitizer and antioxidant effects *in vivo* was unknown. Here, we demonstrated that *A.chilense* oleoresin includes ligands that are able to activate PPARγ with a similar level of activation to that observed with the partial PPARγ activators FMOC-Leu and INT131. In addition, we found a beneficial effect of *Gracilex®* on insulin sensitization in a mouse model of HFD-induced metabolic disorder with a lack of some of the side effects described for the full agonist RGZ on adipose differentiation.

PPARγ is a lipid sensor of the nutritional and metabolic status of organisms (90). As a lipid sensor, this molecule is activated by structurally diverse lipophilic substances because the PPARγ ligand-binding pocket is the largest among the nuclear receptors, allowing the entry and binding of up to two different molecules in a covalent or noncovalent manner (93–96). Our first goal was to evaluate whether the lipids present in *A. chilense* were able to activate PPARγ, comparing it with the full agonist rosiglitazone and with two synthetic SPPARMs, FMOC-LEU and INT131(97, 98). The magnitude of the transcriptional induction of PPARγ achieved by *Gracilex®* was similar to that observed for SPPARM agonists, revealing, as the first finding, that *Gracilex®* possesses SPPARM-like ligands. The fact that similar induction of PPARγ transcriptional activity was induced in two different cell lines (Fig. 1) together with a dose-dependent response suggests that the main bioactive molecules are already present in the extract and that further modifications may not be undergone by intrinsic cell metabolism. Natural extracts presenting PPARγ agonist activity have been reported and reviewed extensively by Wang et al. (50). In a broad screening of extracts derived from 71 different plant species, almost 40% (28 extracts) presented PPARα or PPARγ activity. However, whether this activity possesses insulin sensitizer activity or antioxidant activity *in vivo* was not evaluated. In agreement with our findings, a greater biological activity was found in the extracts produced with dichloromethane than in the extracts obtained with methanol (92), suggesting that the putative PPAR ligands are more easily extracted with apolar solvents. A similar screening was performed by Kim and colleagues (99) using different macroalgae and methanol/ethyl acetate extractions. They found that the extract from *Sargassum yezoense*, a brown seaweed, showed the strongest potency for PPARγ/a transcriptional response in a PPRE-based transcriptional assay. According to their studies, the increased PPAR activity was explained by the presence of sarquinoic acids and sargahydroquinoic acids that could increase adipocyte differentiation of 3T3-L1 cells, which is an unwanted side effect for natural-derived ligands expected to be used in diabetes, since this feature is associated with weight gain in T2DM patients (100). *Gracilex®*, not only induced partial PPARγ activation similar to SPPARMs but also improved insulin sensitivity and normalize glucose levels with no alterations in weight gain and did not promote adipocyte differentiation in a classical *in vitro* model assay. Previously, the content of eicosanoids derived from ARA was studied in *A. chilense* and the closely related *A. vermiculophyllum. A. chilense* contained 8-hydroxyeicosatetraenoic acid (8-HETE) and 7,8-di-hydroxyeicosatetraenoic acid (7,8-diHETE), while *A. vermiculophyllum* contained, in addition to 8-HETE and 7-8-HETE, the prostaglandin PGE2, which was not present in *A. chilense*. This finding suggests that closely related algae can have similar contents of PUFAs and other lipids, but different enzymes associated with their modification. The authors also found that it was possible to promote oxylipin biosynthesis by subjecting the algal tissue to a mechanical damage. Physiologically, these oxylipins are produced after the release of ARA by phospholipase A and conversion by lipoxygenase as defense signals against epiphytes (19,20,22,24). Our procedure for obtaining *Gracilex®* included mechanical trituration of algae to activate its intrinsic defense mechanism and subsequent extraction with DCM, suggesting that this step may increase the oxylipin content of the extract. Further studies will determine if this is the case in the future. A recent analysis of the *A. chilensés* lipids by Honda et al. (19) revealed that glycerolipids comprise the largest proportion of the total lipid contents (monogalactosyldiacylglycerol and phosphatidylcholine), carrying mostly a combination of ARA and palmitic acids (20:4n-6/16:0) in their structure. In agreement with this study, our analysis of the fatty acid profile of *Gracilex®* showed that palmitic acid and ARA were the most abundant fatty acids, representing 40% and 21.1% of the total fatty acids, respectively. Our findings also agree with a recent report on the fatty acid profile of the related red macroalga *Gracilaria* spp. (18). The involvement of these fatty acids and metabolites in the ability of *Gracilex®* to induce PPARγ activation is currently unknown and under investigation in our laboratory. However, it is known that among the endogenous PPARγ ligands described in the literature, 8-HETE is characterized as a dual PPARα and PPARγ agonist (28, 37), whereas the activity of 7,8-di-HETE has not been evaluated. Other lipid compounds present in *Gracilex®* may also act as agonists in a concerted manner, given the ability of the PPAR ligand binding pocket to bind more than one lipid agonist at the same time (26, 101). PUFAs, such as ARA and docosahexaenoic acids (DHA), and their derivative eicosanoids both bind to PPARγ, but eicosanoids are stronger agonists than PUFAs (28,102,103). Other lipids, including lysophosphatidic acids (1-O-oleyllysophosphatidic acid) (104, 105) and phytanic acid (106), also possess PPARγ transactivation activity.

One of the central pathologies in metabolic syndrome is insulin resistance, an event that occurs before T2DM. The development of this pathology directly correlates with overweight and obesity; indeed, obesity explains 60-80% of diabetes cases (107, 108). Here, we provided evidence that *Gracilex®* treatment might benefit overweight patients with insulin resistance. Oral treatment of a mouse model of diet-induced obesity with *Gracilex®* improved insulin and glucose levels, which were altered by the diet. Indeed, the HFD-fed mice treated with *Gracilex®* for one month showed a notable decrease in both blood parameters and glucose and insulin levels at the 300 mg/kg dose, similar to RGZ at 5 mg/kg. RGZ increased the levels of adiponectin, as expected (109). *Gracilex®* significantly decreased the levels of adiponectin in the HFD-fed mice compared to the HFD-fed control mice. This finding indicates that the beneficial effect of *Gracilex®* is not due to the reported capacity of PPARγ agonists to increase adiponectin levels (109). Adiponectin is a hormone that is secreted by adipose tissue and has autocrine and endocrine actions with reported global insulin sensitizer effects (60). Nonetheless, adiponectin increases the growth of fat tissue, and overexpression of adiponectin in 3T3-L1 cells increases lipid storage and adipogenesis (110). Thus, the limited capacity of *Gracilex®* to increase adipocyte differentiation and accumulation of triglycerides measured by oil red O staining in 3T3-L1 cells correlates with the decreased adiponectin levels *in vivo* induced by *Gracilex®* in the HFD-fed mice. Further experiments studying the genetic programs and transcriptomes in 3T3-L1 cells and adipose tissue from HFD-fed mice treated with vehicle and *Gracilex®* will help elucidate this issue since the *adiponectin* gene possesses PPRE and its transcription is regulated by PPARγ (111). The regulation of adiponectin expression is tightly controlled by a number of transcription factors, including C/EBPalpha (CCAAT/enhancer-binding protein alpha), SREBP-1c (sterol regulatory element binding protein) and CREB (cAMP response element binding protein) (112). Therefore, it is possible that lipids contained in *Gracilex®* counteract the signal pathways leading to activation of these transcription factors or that the set of coactivators recruited by the potential PPARγ ligands present in *Gracilex®* represses adiponectin transcription since SPPARMs differentially regulate gene expression compared to TZDs (47, 50).

Another mechanism for the PPARγ effect on insulin and glucose metabolism in addition to increased adiponectin levels has been described. PPARs also generate an increase in the expression of liver enzymes regulating glycogen accumulation in the liver, which reduces the levels of glucose in the blood (113, 114). Obesity downregulates PPARγ transcriptional events that are restored by agonists, such as TZD treatment, including the promotion of adipose tissue angiogenesis and the anti-inflammatory effects on adipose tissue, decreasing systemic inflammation induced by obesity (42).

As a source of multiple types of chemical molecules, *A. chilense* oleoresin showed antioxidant activity, as has been previously reported in diverse algae-derived extracts. While aqueous extracts have been characterized in depth either for their *in vivo* activity or for their antioxidant components (such as phenols and polyphenols, amino acid-like micosporines and sulfated polysaccharides), scarce information exists for the apolar organic-produced fraction like the one we have produced in this study. In the red seaweed *Gracilaria blodgetti,* a chloroform extract could decrease the oxidative stress induced in leukocytes (115–118). These results indicate that apolar extracts can be a source of antioxidants for biological membranes.

Depending on the type of alga, the solvent used and seasonally of algae harvested, a majority or minority of antioxidant components and their activities can be redistributed between polar or apolar fractions (13, 119). Difficulties in determining the *in vitro* capacity of apolar extracts or molecules through classical methods are known, and more than one assay should be used to confirm their antioxidant potential (120, 121). Considering the few alternative methods for determining antioxidant capacity in oleoresins, in addition to the *in vitro* TAC method, we evaluated the *in vivo* antioxidant capacity of *Gracilex®* using a strain mutant of *C. elegans* that is sensitive to oxidative stress (72). We found that nematodes fed *Gracilex®* were protected against exposure to hydrogen peroxide with a similar magnitude as that given by N-acetyl-cysteine (NAC) treatment, reflected by an increase in nematode survival. NAC is a source for the production of glutathione, and it has been shown to prevent ROS generation (89) and to increase oxidative stress protection in *C. elegans* (88). Therefore, it is an appropriate positive control for our experiments. These results correlated with high contents of tocopherols and ß-carotene in *Gracilex®*, which are well-known antioxidants reducing lipid peroxidation (122–124).

## 5. Conclusion

Altogether, our results indicate that *Gracilex®*, an oleoresin produced from *A. chilense* after dicloromethanol extraction, contains natural PPARγ ligands that increase its transcriptional activity, with insulin sensitizing and antioxidant properties *in vivo*.

## Author contribution

CP, KF and FCB performed the conceptualization of the study. CP conducted the mice studies. The cell studies were performed by CP, GL, CG, YS and LL. Preparation of the *A.chilense* oleoresin was performed by CP, GL, VB, LL, RI. Availability of *A. chilense* biomass was possible by FC and LCP. VB performed CUPRAC antioxidant determination. RI and RA performed *C.elegans* studies. Supervision of experiments was performed by KF, RA, LCP and FCB. KF and FCB performed original draft preparation. Review and editing were the work of FCB, KF, LCP and RA.

## Funding

Agencia Nacional de Investigación y Desarrollo (ANID) project FONDEF ID16I10185 (FCB) and Basal Center of Excellence in Science and Technology (PIA BASAL AFB170005) (FCB). CORFO “Consorcio tecnológico desarrollo sinérgico de Ingredientes Funcionales y Aditivos Funcionales IFAN” 16PTECAI-66648 (FCB and LCP).

## Conflict of interest

Authors declare no conflict of interest.

## Patents

The methodology to prepare the *A.chilense* oleoresin is fully described in the patent application WO/2014/186913. “Method for the preparation of an oleoresin originating from a red alga that maintains the capacity to induce the transcriptional activity of the nuclear receptor PPAR-Gamma”. PCT/CL2013/000031 Pontificia Universidad Católica de Chile. Inventors: Bronfman Miguel, Bronfman Francisca, Pinto Claudio, Pissani Claudia, Paredes Martínez, María José.

---

**Supplementary Figure 1.**
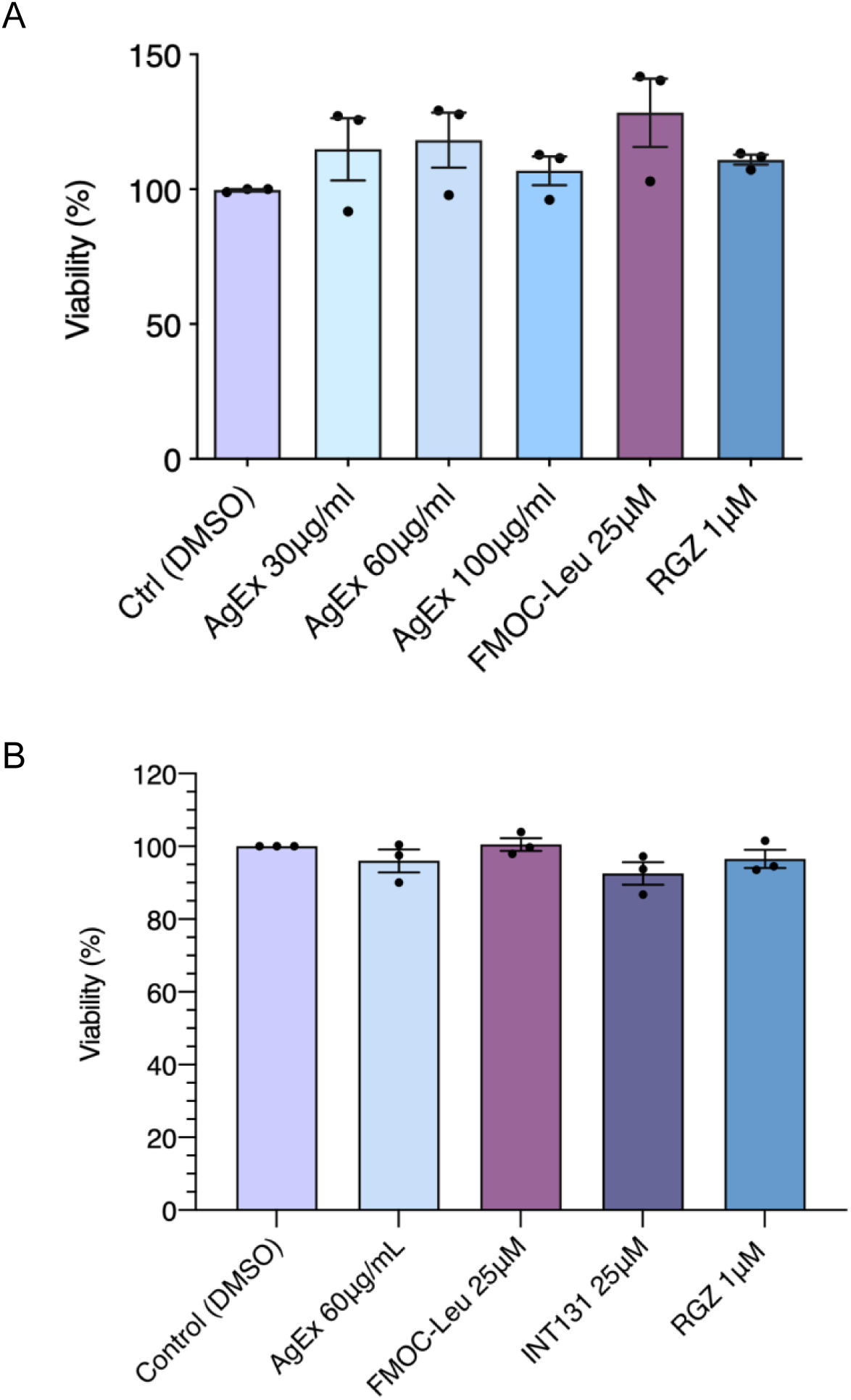
Cell viability of PC12 and 3T3 cells after treatment with PPARγ agonists and *A. chilense* oleoresin. (**A**) PC12 cells were cultured without or with FMOC-Leu, RGZ (1 μM) and *A. chilense oleoresin* or *Gracilex®* (*AgEx)* with three different concentration 30, 60, 90 μg/mL for 24 h. Then, the cells were incubated with MTT for 2 h at 37°C. MTT is used as an indicator of mitochondrial activity and cellular viability. The graph indicates viability relative to the control condition (treatment with vehicle, DMSO). No significant differences were found between treatments with the PPARγ agonist and *AgEx* compared to the control (one-way ANOVA, Tukey’s post hoc test). Values are expressed as the mean (n=3) ± SEM. (**B**) The 3T3-L1 preadipocytes were cultured without or with FMOC-Leu and INT131 at a concentration of 25 μM, rosiglitazone (RGZ, 1 μM) and *AgEx* (60 μg/mL), and at the end of the differentiation period, the cells were incubated with MTT for 2 h at 37°C. The graph indicates viability relative to the control condition (treatment with vehicle, DMSO). No significant differences were found between treatments with PPARγ agonists and *AgEx* (one-way ANOVA, Tukey’s post hoc test, n=3).

**Supplementary Figure 2.**
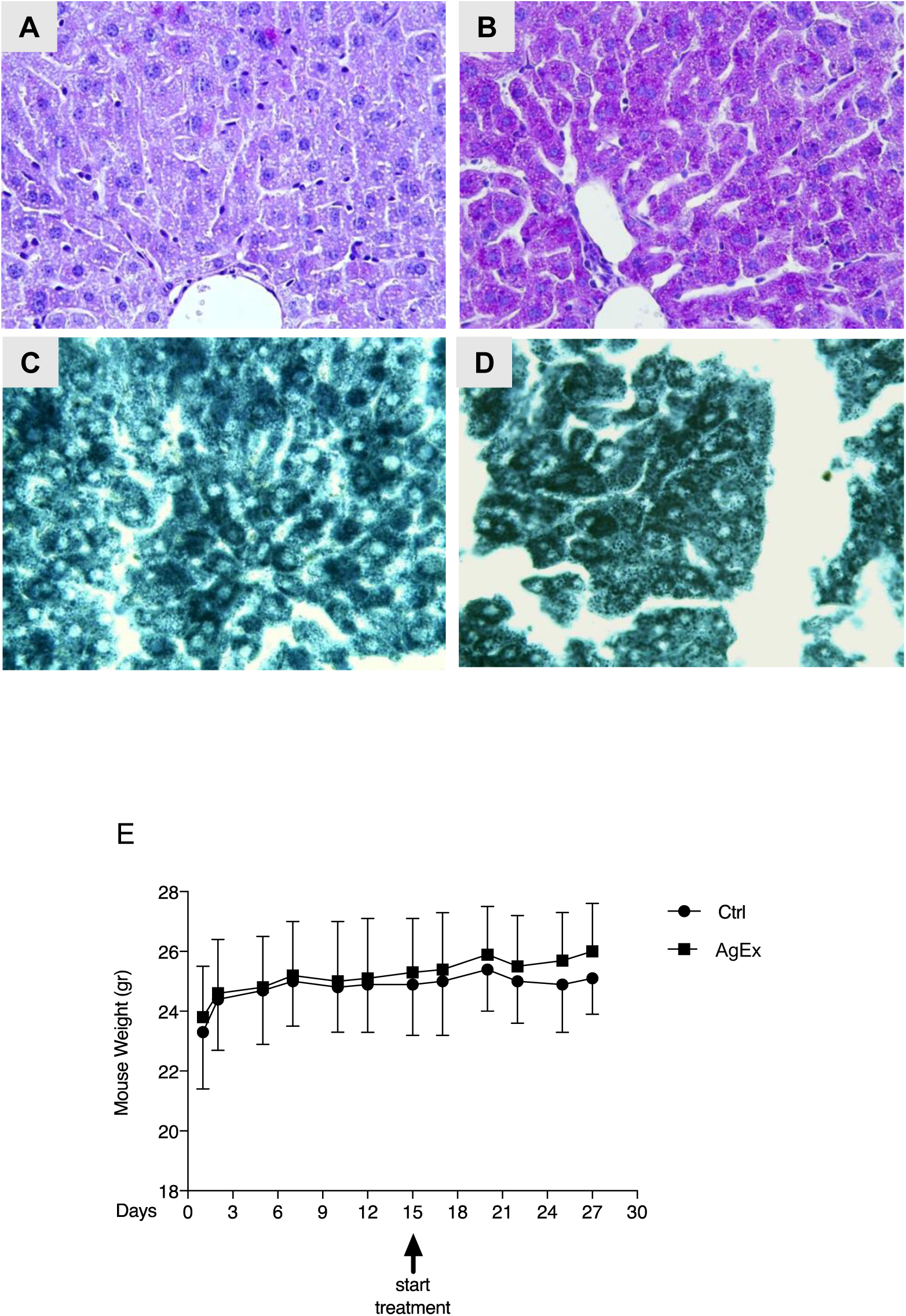
Histological analysis of liver and weight measurements after chronic treatment with *A. chilense* oleoresin in mice fed a normal chow diet. Three-month-old C57BL/6J mice were treated daily for 15 days with 300 (mg/kg) of *A. chilense* oleoresin or *Gracilex®* (*AgEx)* or corn oil (vehicle) extract. Light microphotograph of liver paraffin sections (400X). Hepatocytes were stained with periodic acidic-Schiff (PAS) staining to evaluate glycogen deposits (**A and B**) or with Sudan Black B staining for lipid accumulation (**C and D**). (**A**) and (**C**) are sections from control mice (vehicle). (**B**) and (**D**) are sections from control mice treated with *AgEx*. Mild to moderate glycogen storage was found in the group treated with *AgEx* compared to corn oil (vehicle). No difference was apparent in Sudan Black B staining between the groups. (**E**) Weight curves of normal chow-fed mice treated with *AgEx* for histological study. Three-month-old C57BL/6J mice were weighted 3 times per week, and after 15 days, daily treatment with *AgEx* 300 (mg/kg) or corn oil (vehicle) for an additional 15 days was started. Mice were weighted 3 times per week. The graph shows the average weight curve per group ± SEM.

## Notes

### Competing Interest Statement

The authors have declared no competing interest.

